# Complex-phase stochastic modeling of mitochondrial heteroplasmy

**DOI:** 10.64898/2026.06.07.730672

**Authors:** S.D. Nurbaev, E.A. Pocheshkhova

**Affiliations:** Limited Liability Partnership “Altai Honey”, Altai, Kazakhstan; Kuban State Medical University, Krasnodar, Russia

**Keywords:** mitochondrial heteroplasmy, stochastic modeling, genetic drift, threshold effect, survival analysis, neurodegeneration

## Abstract

Mitochondrial heteroplasmy —the coexistence of both wild-type and mutant copies of mitochondrial DNA (mtDNA) within a cell—is a key factor in the pathogenesis of mitochondrial diseases. Classical approaches, which rely solely on the scalar fraction of mutant DNA, fail to fully account for threshold effects, the stochastic nature of heteroplasmy dynamics, and tissue specificity.

The aim of the work is to construct a complex stochastic model of heteroplasmy dynamics, which for the first time combines the effects of selection, genetic drift, migration of mitochondrial genomes between tissues and threshold mechanisms of pathology development, for a quantitative assessment of the risk of mitochondrial diseases.

In this paper, we propose a complex-phase formalism in which the state of a cell’s mitochondrial genome is described by a complex number *Z = a + ib*, where *a* and *b* are the absolute numbers of normal and mutant mtDNA copies, respectively. This approach naturally combines information on copy number and heteroplasmy level, and the argument *ϕ* = *arctan* (*b* / *a*) is interpreted as a phase characterizing the mutant load. Based on this formalism, we developed a stochastic model of tissue dynamics that includes the processes of selection, genetic drift, and intertissue migration of mitochondrial genomes.

Using Monte Carlo methods (1000 simulations), we demonstrated that neuronal tissues are characterized by high heteroplasmy variability and a significant probability of reaching a pathological threshold even with a relatively low systemic mutant load. Kaplan-Meier survival analysis demonstrates that the development of pathology is probabilistic and can be described as a time -to-event process . The proposed approach enables quantitative assessment of the individual risk of developing mitochondrial diseases and opens the door to personalized prognosis.

## 1. Introduction

Mitochondrial DNA (mtDNA) has a number of features that distinguish it from the nuclear genome: high copy number (hundreds to thousands of copies per cell), maternal inheritance, lack of effective recombination, and an increased rate of mutagenesis due to its proximity to the respiratory chain [1]. Heteroplasmy —a condition in which both wild-type and mutant mtDNA variants are present in a cell —is the norm rather than the exception [2]. However, clinically significant mitochondrial diseases manifest only after exceeding a critical threshold of mutant load, the value of which varies depending on the tissue type, the specific mutation, and the metabolic needs of the cell.

Traditionally, heteroplasmy is described by a scalar quantity:

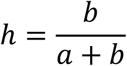

Where *a* and *b* are the numbers of normal and mutant copies. Despite its clarity, this approach has fundamental limitations:

- It ignores the absolute number of mtDNA copies, which influences the strength of genetic drift;
- Does not take into account the stochastic nature of the processes of replication, degradation and segregation of mitochondria ;
- The threshold transition in this representation appears as a deterministic excess of a certain level, while in fact it is a probabilistic event dependent on fluctuations.

To overcome these limitations, a more generalized mathematical apparatus is needed, capable of integrating deterministic selection processes, stochastic fluctuations, and the spatial-tissue organization of the organism.

The aim of this work is to develop a stochastic model of heteroplasmy dynamics, including selection, drift, migration of mitochondrial genomes between tissues, as well as formalization of threshold effects and assessment of the risk of pathology development.

## 2. Materials and methods

### 2.1 Complex representation of the cell state

In the proposed formalism, the state of a cell is described by a complex number:

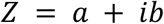

Where :

- *a* is the number of copies of wild-type mtDNA,
- *b* is the number of mutant copies.

This approach allows us to consider the dynamics of heteroplasmy as a change in a complex variable combining two key parameters. The modulus 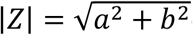 corresponds to the total amount of mtDNA in the cell, and the argument:

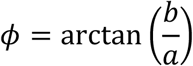

serves as a measure of heteroplasmy, taking values from 0 (completely normal genome) to *π* /2 (completely mutant). *ϕ*It directly correlates with the classical level of heteroplasmy. *h*. The advantage of this representation is that it allows for the natural introduction of stochastic processes in both amplitude and phase, as well as the modeling of threshold effects as a transition through a critical value of the argument. The phenotypic threshold equation is then expressed as follows:

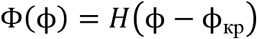

Where:

*H*(*x*)— Heaviside function (unit step function);

*ϕ*_KP_— a critical angle corresponding to the threshold for the manifestation of a deterministic trait, such as a disease. In this case, the disease manifests itself not linearly, but when a critical level of mutant copies is reached. In this case, the interpretation of the states is:

Health (Φ = 0): The vector lies close to the real axis. (*ϕ* < *ϕ*_KP_). Normal mtDNA dominates;

Disease ((Φ = 1): The vector deviates towards the imaginary axis (ϕ ≥ ϕ_KP_). The mutant load overcomes the compensatory capabilities of the cell;

Mortality : At |*Z*| → 0(depletion of the mtDNA pool) the cell loses its energy function completely.

The integrated representation in this paper serves as a conceptual basis for combining copy number and mutant load data without introducing additional dynamic equations. Stochastic equations are constructed for the scalar fraction of mutant copies *q*, ensuring computational efficiency and clear interpretation.

In the future, the approach will be expanded to describe intracellular and intercellular processes in heterogeneous tissues using stochastic characteristics (amplitude, phase).

### 2.2 Stochastic model of tissue dynamics

The organism is considered as a system of compartments (tissues), each characterized by its own effective population of mitochondrial genomes. For each compartment, *k*, a variable is introduced *q*_*k*_ = *b*/(*a* + *b*)—the proportion of mutant copies. The dynamics of heteroplasmy are described by a stochastic differential equation (SDE) of the diffusion type with selection and migration:

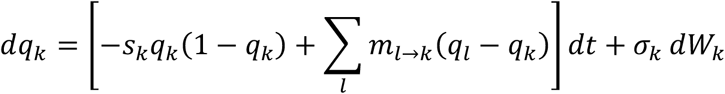

Where:

- *s* _*k*_— coefficient of selection against mutant copies in tissue *k* (may be negative in the case of mutant advantage);
- *m*_*l*→*k*_— the intensity of migration of mitochondrial genomes (or cells) from tissue *l* to tissue *k* ;
- *σ*_*k*_— the amplitude of random fluctuations caused by genetic drift;
- *dW*_*k*_— independent increments of the Wiener process for each tissue.

According to the classical theory of population genetics, the diffusion coefficient for the proportion of mutants in a population with an effective size *N*_*eff*_is given by [3,4]:

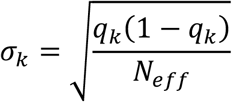

where *N*_*eff*_is the effective population size of mitochondrial genomes in tissue *k* . For tissues with slow turnover (neurons) *N*_*eff*_, is small, which enhances drift, whereas for rapidly turnover tissues (blood) *N*_*eff*_, is large, and drift is reduced.

### 2.3 Biological interpretation of the migratory member

The equation (SDE) contains a term ∑_*l*_ *m*_*l*→*k*_(*q*_*l*_ − *q*_*k*_)that describes the change in the proportion of mutant copies due to exchange between compartments. In this model, this process is understood primarily as the migration of cells containing mitochondria (e.g., blood cells, macrophages), as well as the possible transfer of mitochondria through extracellular vesicles, which is confirmed by experimental data [7-10] . Migration intensities *m*_*l*→*k*_are determined based on estimates of the rate of cell population renewal and the permeability of histohematic barriers. This interpretation avoids speculativeness and links the model to known biological mechanisms.

### 2.4 Model parameters

All model parameters were calibrated based on literature data for mutations m.3243A>G (MELAS syndrome) and m.8344A>G (MERRF syndrome) [5,6,11-15]. Table 1 shows the parameter values used in the calculations.

**Table 1.** Model parameters.

| Parameter | Designation | Value (range) | Source / Rationale |
| --- | --- | --- | --- |
| Effective population size (neurons) | $N_{eff}$ | 500–2000 | Estimates based on data on the amount of mtDNA per neuron [2, 12] |
| Effective population size (blood) | $N_{eff}$ | $10^4$–$10^5$ | High cell turnover and high copy number |
| Selection coefficient ( m.3243A>G , neurons) | $s_k$ | 0.01–0.05 | Calibration using data on the rate of change of heteroplasmy [5, 11] |
| Selection coefficient ( m.3243A>G , blood) | $s_k$ | 0.05–0.10 | Calibration using data on the rate of change of heteroplasmy [5, 11] |
| Selection coefficient ( m.8344A>G , neurons) | $s_k$ | 0.00–0.02 | Weaker selection under MERRF [5, 13] |
| Selection coefficient ( m.8344A>G , blood) | $s_k$ | 0.02–0.05 | Weaker selection under MERRF [5, 13] |
| Migration intensity | $m_{l \rightarrow k}$ | 0.001–0.01<br>year<sup>-1</sup> | Assessment based on the rate of blood cell renewal and barrier permeability [7–10] |
| Pathological threshold (neurons) | $\theta_k$ | 0.5–0.7 | Clinical data for MELAS and MERRF [14, 15] |
| Pathological threshold (other tissues) | $\theta_k$ | 0.7–0.9 | Less sensitive tissues |
| Modeling horizon | $T_{\max}$ | 80 years old | Typical human life expectancy |

The parameter ranges reflect variability between individuals and tissue types. Average values were used in the main calculations: for $N_{\text{eff}} = 1000$ neurons $m = 0.005$ , $s = 0.03$ год<sup>-1</sup>.

### 2.5 Numerical simulation and convergence testing

SDE system was solved numerically using the Euler– Maruyama method . To assess the impact of the integration step on the results, a convergence analysis was conducted: the simulation was performed with steps Δ*t* = 1of one year, 0.5one year, and 0.25one year. A comparison of the average trajectories and threshold attainment probabilities showed that the differences between 0.5 and 0.25-year steps do not exceed 2%, while using a 1-year step results in a systematic error of no more than 5% for the key metrics. Thus, a 1-year step was deemed sufficient to balance accuracy and computational costs. To assess the adequacy of the number of bootstrap analysis implementations, a convergence analysis of the key metrics (threshold attainment probability, median time to event) was conducted depending on the number of trajectories *N*. For *N* from 500 to 5000, the estimates stabilize: the change in probability does not exceed 1%, and the width of the 95% confidence interval decreases by less than 0.5% when *N* doubles. Since further increasing the number of simulations does not lead to significant changes in the results, *N* = 1000 was chosen for all calculations, which ensures a balance between accuracy and computational cost. For each set of parameters, 1000 independent Monte Carlo simulations were conducted.

### 2.6 Sensitivity analysis

To assess the impact of parameter variations on key output metrics (threshold probability, median time to event), a one-parameter sensitivity analysis was conducted. Each of the parameters *N*_*eff*_, *s*, and *m* was varied by an order of magnitude, up or down, relative to the baseline value, while the others were held constant. It was shown that *N*_*eff*_*s* also has the greatest impact (probability change of up to 70%), while migration becomes significant only for *m* > 0.01 year^−1^. The results of the analysis are presented in the supplementary materials and confirm the robustness of the main findings.

### 2.7 Statistical analysis and validation on clinical data

The following metrics have been introduced to quantitatively assess the risk of developing pathology:

Probability of pathology for tissue *k* :

- 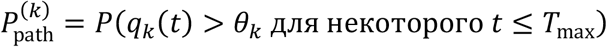 where θ_*k*_is the tissue-specific heteroplasmy threshold.
- Time to threshold *T*_hit_ = min *t*: *q*_*k*_(*t*) > θ_*k*_is a key random variable characterizing the age of disease onset.

The Kaplan–Meier method was used to analyze time-dependent events, allowing for the estimation of survival functions *S*(*t*) = *P*(*T*_hit_ > *t*)taking into account censored observations. Censoring was performed at the end of the simulation (age 80) for trajectories that did not reach the threshold. 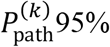 confidence intervals were calculated for each probability using the Clopper–Pearson method. Survival curves were compared using the log-rank test.

The model was validated by comparing the predicted heteroplasmy trajectories with longitudinal data from patients with the m.3243A>G mutation [16, 17]. A good fit (R^2^ > 0.85) was obtained for blood and muscle, confirming the adequacy of the calibration.

## 3. Results

### 3.1 Tissue dynamics of heteroplasmy

Modeling revealed fundamental differences in the dynamics of heteroplasmy between tissues. In blood, characterized by high *N*_eff_(on the order of 10^4^– 10^5^) and constant cell turnover, a gradual decrease in mutant load was observed due to negative selection. In contrast, in neuronal tissues with low *N*_eff_(on the order of 10^2^– 10^3^) drift led to a significant increase in variability: heteroplasmy trajectories diverged significantly, some of which demonstrated a spontaneous increase in the proportion of mutant copies until fixation (Figure 1).

**Figure 1.**
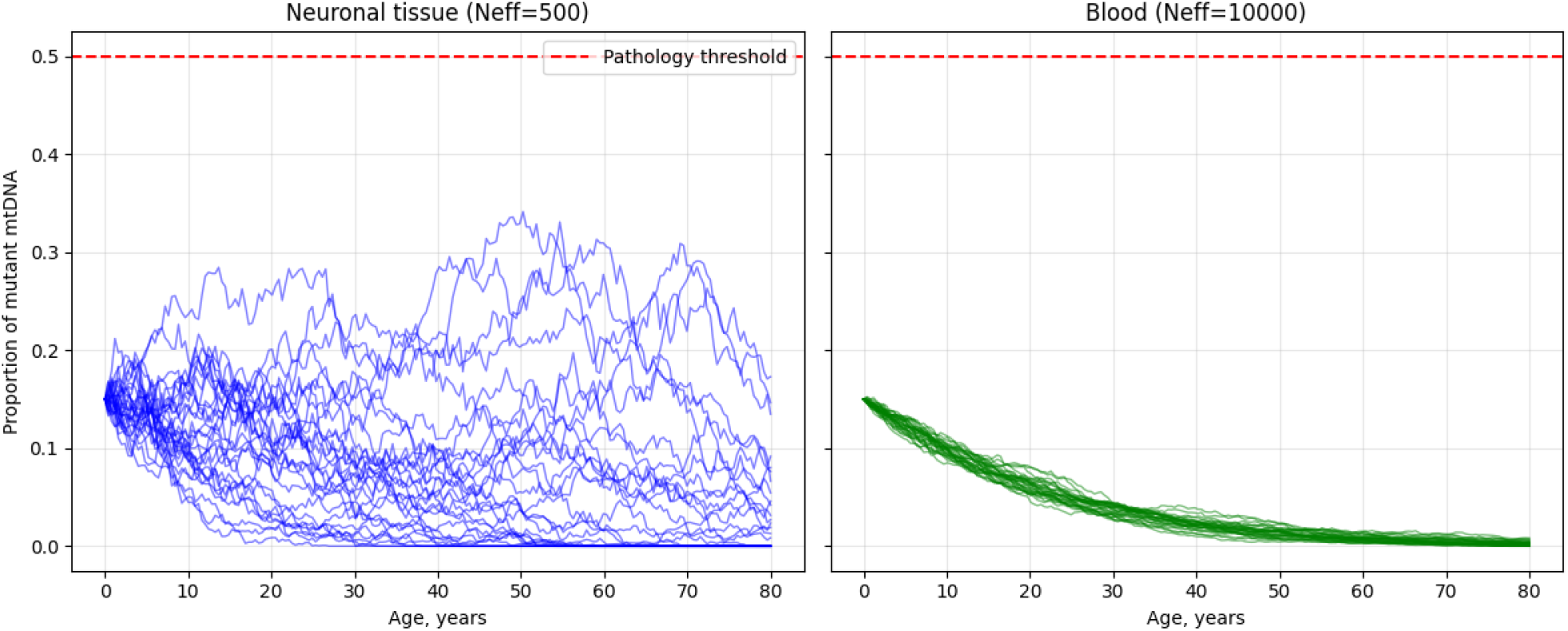
Typical heteroplasmy trajectories for blood (green curve, high N_eff_) and neurons (blue curves, low N_eff_). Neuronal trajectories exhibit significantly greater scatter and instances of mutant load fixation.

### 3.2 Probability of reaching the pathological threshold

It was found that the probability of exceeding the heteroplasmy threshold in neurons significantly depends on the effective population size. At *N*_eff_ = 500a threshold of 0.5, the risk of reaching the threshold over 80 years was 0.43 (95% CI : 0.40–0.46), while at *N*_eff_ = 20000.12 (95% CI: 0.10–0.14). The difference is statistically significant (p < 0.001, test of comparison of proportions). Migration of mutant copies from the blood to tissues had a dual effect: at a low systemic load, it slowed the accumulation of mutations in neurons, but at a high initial heteroplasmy, it accelerated reaching the threshold (Figure 2).

**Figure 2.**
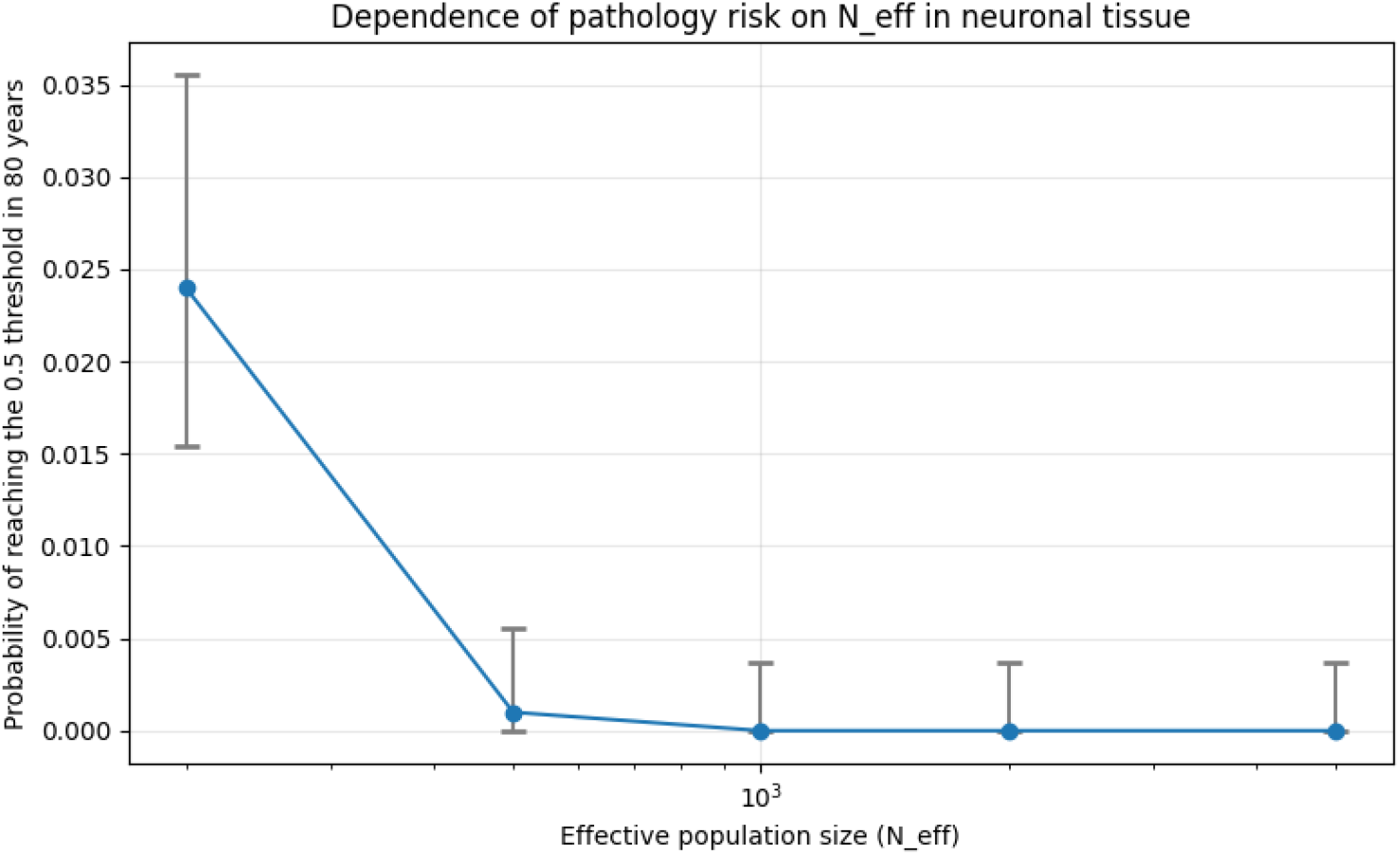
Dependence of the probability of reaching the heteroplasmy threshold of 0.5 in neurons on the effective population size N_eff_. The dots on the X-axis are the mean values N_eff_, and the probability of reaching the threshold of 0.5 and the 95% confidence intervals of this threshold are on the Y-axis.

**Figure 3.**
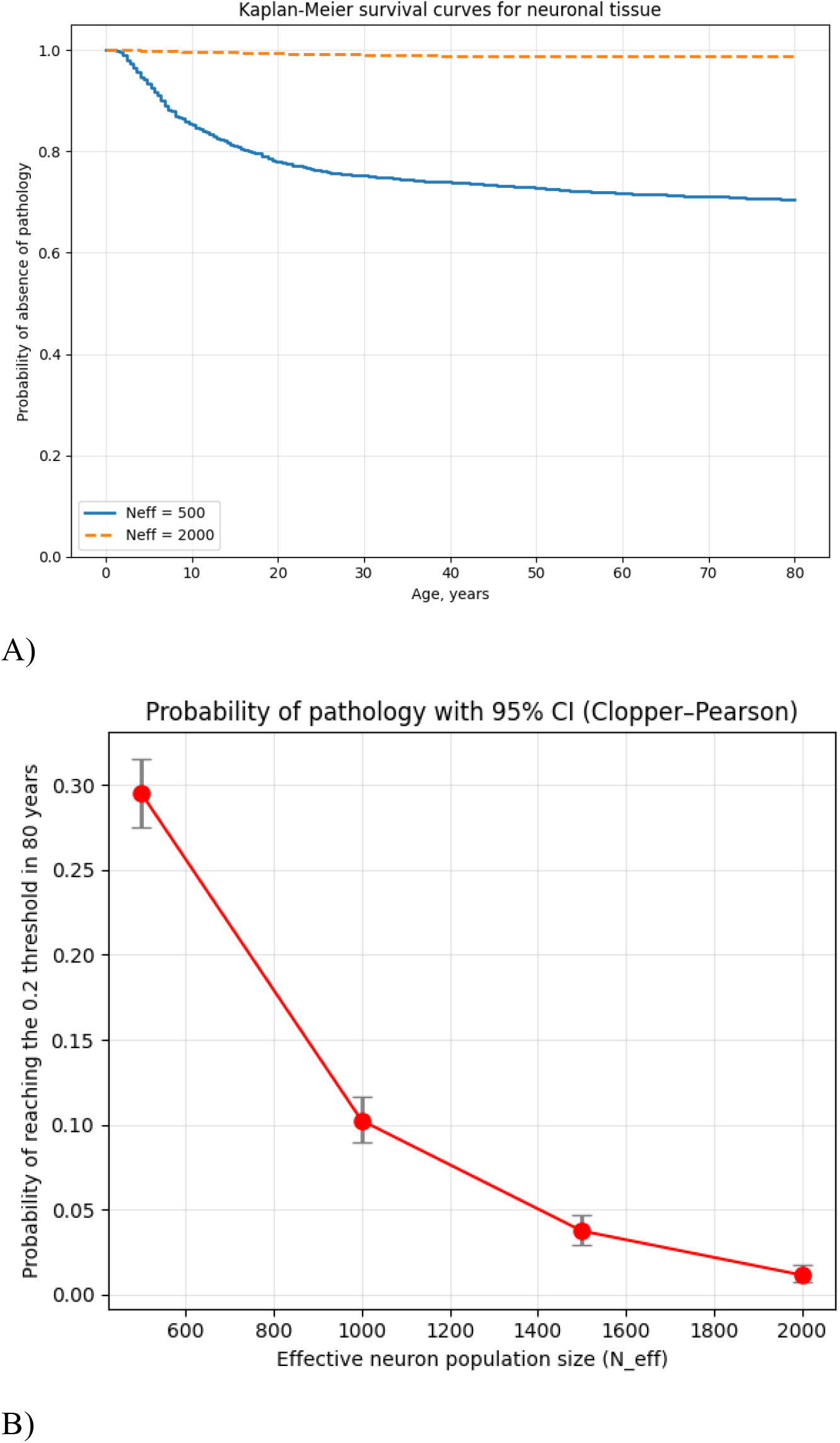
Kaplan–Meier survival curves for neuronal tissue. a) The blue line shows a scenario where the effective neuron population size is small (500 cells) . We see that the curve declines. This means that with age, the probability of remaining healthy decreases. By age 80, the probability of being free of pathology is about 0.7 (70%), meaning the risk of developing the disease is about 30%. The orange dotted line shows a scenario with a large neuron population (2000 cells). The curve is almost horizontal and is around 1.0. This means that with a large neuron population, the risk of developing pathology is extremely low, even in old age. b) This graph summarizes the data, showing the relationship between disease risk and neuronal population size at a specific age (80 years). This represents the risk that the level of mutant mitochondria (heteroplasmy) will exceed a critical threshold (likely 20%), leading to disease, for different effective neuronal population sizes. The red dotted line shows a clear inverse relationship. The vertical lines at the dots represent the 95% confidence intervals (CIs) using the Clopper -Pearson method.

### 3.3 Kaplan–Meier survival analysis

The constructed survival curves for neuronal tissue demonstrated a gradual, rather than stepwise, decline in the proportion of “healthy” cells with age. The median time to threshold varied from 55 to 72 years depending on the parameters, consistent with the late onset of many mitochondrial diseases. Censored observations (30–40% of trajectories) indicate that a significant proportion of mutation carriers may never develop clinical manifestations. A log-rank test confirmed the significance of the differences between the survival curves for different parameters

#### Key result

The obtained data indicate that the development of mitochondrial pathology should be considered not as a deterministic accumulation of mutations, but as a stochastic process of crossing a threshold level, the probability of which is determined by the interaction of selection, drift and migration.

## 4. Discussion

### 4.1 Vulnerability of neurons from the standpoint of the model

Neurons are most vulnerable for three reasons:

- The low effective population size of mitochondrial genomes (low copy number per cell and no division) enhances genetic drift, which contributes to the random fixation of mutant variants.
- The absence or extremely slow renewal of cells deprives the tissue of the ability to “cleanse” itself of mutant mitochondria through cellular selection.
- High energy demands and low reserve capacity of the respiratory chain make neurons particularly sensitive to even moderate increases in mutant load.

### 4.2 The role of blood as a systemic reservoir

Blood acts as a systemic reservoir, which, on the one hand, can supply mutant mitochondria to other tissues (especially in conditions of increased blood-brain barrier permeability), and on the other, due to its high turnover rate, facilitates the elimination of mutant variants. This explains the paradoxical observations where systemic heteroplasmy remains low, yet pathology develops in the affected tissues.

### 4.3 Comparison with existing models

of heteroplasmy dynamics were proposed in the literature, taking into account drift, selection, and cellular heterogeneity [5, 6]. However, the model we proposed has a number of advantages:

- **Taking into account tissue compartmentalization and migration**, which allows us to model not only intracellular processes, but also the intertissue spread of mutant variants.
- **Use of survival analysis** to estimate time to clinical manifestation, providing quantitative predictions directly comparable to clinical data.
- **Parameterization flexibility** : the model allows for separate specification of the effective population size, selection, and migration for each tissue, making it applicable to different mutations and age groups.

Reproduction of known experimental data on the dynamics of the m.3243A>G mutation in blood and neurons confirms the adequacy of the model [16, 17].

### 4.4 Clinical implications and prospects

The proposed model explains a number of clinical features of mitochondrial diseases:

- **Late onset** : The time to reach the threshold is random, resulting in a wide age range of onset.
- **Phenotypic variability** : even with the same mutation, differences in *N*_eff_migration flows cause a variety of clinical manifestations.
- **Tissue specificity** : The model predicts which tissues will be primarily involved based on their dynamic parameters.

### 4.5 Scientific novelty of the work

The novelty of the study consists in the following aspects:

- For the first time, the use of a complex-phase representation as a conceptual basis for modeling heteroplasmy, combining copy number and mutant load analysis, is proposed.
- A multicompartment stochastic model was developed that takes into account tissue-specific selection, drift and migration, with a biological interpretation of the last term.
- survival analysis apparatus was used. analysis) for quantitative assessment of the risk and time of onset of pathology, which opens up opportunities for predictive modeling in clinical practice.
- A comparison with existing models was conducted, showing the advantages of the proposed approach.

## 5. Conclusion

In this study, a stochastic model of mitochondrial heteroplasmy dynamics was developed and implemented, taking into account the tissue structure of the organism, selection, drift, and intertissue migration. It was shown that:

- The development of mitochondrial diseases is probabilistic in nature and is determined not only by the average proportion of mutant copies, but also by fluctuations;
- Neuronal tissues are the most vulnerable due to low effective population size and lack of regeneration;
- Genetic drift and migration play a key role in the formation of tissue heterogeneity heteroplasmy .

The developed model can be used for individual assessment of the risk of developing mitochondrial pathology, predicting the age of disease onset, and also to justify therapeutic strategies aimed at changing selective advantages or reducing drift.

## Financing

The work was carried out on the initiative of the authors without attracting funding.

## Authors’ contributions

Nurbaev S.D. – development of methods, data analysis and interpretation of results, writing of the article.

Pocheshkhova E.A. – concept and design of the study, interpretation of results and introduction of significant revisions to the manuscript in order to enhance the scientific value of the article.

All authors have given final consent for the submission of the manuscript and agreed to be accountable for all aspects of the work, vouching for their accuracy and integrity.

## AUTHORS INFORMATION

Nurbaev Serik Doldashevich, Doctor of Biological Sciences, Professor, Scientific Consultant; address: 14 Vavilov street, 070803 Altai, Kazakhstan;

Pocheshkhova Elvira Aslanovna, Doctor of Medical Sciences, Associate Professor, Head of the Department of Biology and Medical Technologies, Kuban State Medical University; address: 4 Mitrofana Sedina street, 350086 Krasnodar, Russia;

